# Impact of immediate cryopreservation on the establishment of patient derived xenografts from head and neck cancer patients

**DOI:** 10.1101/2020.02.03.930891

**Authors:** Lindsey Abel, Arda Durmaz, Rong Hu, Colin Longhurst, Andrew M. Baschnagel, Deric Wheeler, Jacob G. Scott, Randall J. Kimple

## Abstract

**Background:** Patient-derived xenografts established from human cancers are important tools for investigating novel anti-cancer therapies. Establishing PDXs requires a significant investment and many PDXs may be used infrequently due to their similarity to existing models, their growth rate, or the lack of relevant mutations. We performed this study to determine whether we could efficiently establish PDXs after cryopreservation to allow molecular profiling to be completed prior to implanting the human cancer.

**Methods:** Fresh tumor was split with half used to establish a PDX immediately and half cryopreserved for later implantation. Resulting tumors were assessed histologically and tumors established from fresh or cryopreserved tissues compared as to the growth rate, extent of tumor necrosis, mitotic activity, keratinization, and grade. All PDXs were subjected to short tandem repeat testing to confirm identity and assess similarity between methods.

**Results:** Tumor growth was seen in 70% of implanted cases. No growth in either condition was seen in 30% of tumors. One developed a SCC from the immediate implant but a lymphoproliferative mass without SCC from the cryopreserved specimen. No difference in growth rate was seen. No difference between histologic parameters was seen between the two approaches.

**Conclusions:** Fresh human cancer tissue can be immediately cryopreserved and later thawed and implanted to establish PDXs. This resource saving approach allows for tumor profiling prior to implantation into animals thus maximizing the probability that the tumor will be utilized for future research.

## Introduction

Patient derived xenografts (PDXs) are important tools for investigating novel anti-cancer therapies, elucidating mechanisms of oncogenesis and therapeutic response, and understanding drivers of therapeutic resistance and tumor evolution. PDXs represent a valuable resource for pre-clinical translational oncology as they allow investigators to sample the heterogeneity within a population of cancer patients. When properly managed, PDXs are a renewable resource that can be made available through biobanking for drug screening, therapeutic development, mechanistic confirmation, and basic science discovery (1,2).

The process of generating PDX involves a highly coordinated effort on the part of multiple entities. Individuals must work together to collect time and temperature sensitive samples while complying with federal and state regulations to protect patients and their personal data. The most common approach to establishing PDXs, and the one used at our institution, involves obtaining fresh tissue and as quickly as possible implanting the tissue into recipient mice (3,4). We have previously demonstrated that this implantation can occur up to 48 hours after donation, if the tissue is appropriately stored (4). Resulting PDXs can be cryopreserved or passaged for experiments or expansion but are typically used within five to ten passages. Initial tumor implantation and growth often occurs at the same time as detailed genomic or transcriptomic analysis thus resulting in significant costs (mice, cage charges) dedicated to the generation of PDXs which may find little subsequent use due to duplication of specific subtypes that have been previously established.

We performed this study to provide evidence that we can immediately cryopreserve tumor specimens and later thaw them to use for PDX establishment rather than immediately implanting all tissue specimens. In cases where sufficient tumor was present to enable both immediate implantation and cryopreservation, we compared PDX establishment success rates and tumor histology between the two approaches. It is our hope that this evidence will enable investigators to maximize resources by only establishing PDXs that meet specific criteria necessary for future experiments by allowing time for tissue characterization (i.e. specific cancer diagnosis, mutational profile, tumor biomarkers) prior to implantation and expansion in animals.

## Materials and Methods

### Receiving and Processing primary (P0) tumor tissue

We have previously described our approach to establishing PDXs (3,4). Briefly, fresh tumor tissue was obtained from patients consented for deidentified excess tissue donation under an IRB approved protocol via the institutional Translational Sciences Biobank (IRB #UW-2016-0934, expiration 8/11/2020). PDXs were established under an IRB-exempt protocol utilizing this de-identified, residual tissue (IRB exemption #2016-0570). Tissue was stored for less than 48 hours in DMEM (catalog number 10-013-CV) at 4°C prior to transfer to the investigational team. In cases in which at least 0.6 g of tissue were provided, samples were processed for engraftment and cryopreservation. Briefly, tissue was rinsed and cleared of blood and/or necrotic tissue, placed into 400 μl of prepared PDX media, and minced with sharp, sterile scissors to obtain a slurry. This slurry was divided equally: half used for implantation into animals (i.e. fresh) and half for cryopreservation. PDX media and phosphate buffered saline was prepared fresh per recipe as previously described (3–5).

### Tissue Cryopreservation

Minced tissue was pipetted from the Eppendorf tube and placed into a cryovial. PDX media was used to bring the total volume to 540 μl prior to addition of 60 μl of DMSO (final volume of 600 μl). The sample was mixed by repeated gentle manual pipetting (10 times). Tissue was placed in a controlled rate freezer container (Bel-Art Cat #F18844-0000) filled according to manufacturer instructions with room temperature isopropanol. The entire container was placed in a –80°C freezer for 24 hours prior to transfer of cryovials to a vapor phase liquid nitrogen freezer for long-term storage at −148°C.

### Tissue reanimation

Cryopreserved patient tissue was removed from its liquid nitrogen storage and placed in a warm bead bath at 37°C. Immediately after tissue was thawed, it was pipetted from the cryovial into a 1.5ml Eppendorf tube filled with 600 μl of 1X PBS. The mixture was pelleted by centrifugation at 200 × g for 2 minutes at 4°C. The supernatant was removed, the cell pellet resuspended in PBS, and the pelleting step repeated. Depending on the amount of tissue and the number of sites to be engrafted, the cell pellet was resuspended with 200-400 μl of PDX media.

### Tissue engraftment to SCID mice

Tumor engraftment was performed as previously described (3,4). All animal studies were performed under an IACUC approved protocol (UW IACUC #M005974) in accordance with standards for ethical animal care. Briefly, 4-6 week old male and female NOD-SCID gamma (NSG, NOD.Cg-*Prkdc^scid^ Il2rg^tm1Wjl^/SzJ*) mice have prepared tumor tissue injected subcutaneously into the flank. Tumor in PDX media is mixed in a 1:1 ratio with Matrigel (Corning, cat #CB 40234C) by gentle pipetting and stored on ice to prevent polymerization. Animals are anesthetized using isoflurane, tumor tissue is implanted using a 18g needle using 100-200 μl of tumor-Matrigel slurry. Following removal of the needle, the site was reapproximated using a gentle pinch for 10-20 seconds to minimize leakage. Animals were returned to their cages and monitored until they had recovered from anesthesia.

Mice were monitored weekly until tumors reached a size of at least 500mm^3^. To harvest tissue for analysis, mice were euthanized using CO_2_ and cervical dislocation. Tumors were collected and divided for multiple uses: formalin fixation and paraffin embedding, flash frozen tissue, and cryopreservation. The remaining tissue was engrafted into another SCID mouse for successive passages.

### Histologic analysis

A fraction of retrieved tumor from the NSG mice was fixed by 10% formalin and processed for routine paraffin embedding. Paraffin embedded sections (5 μm) were stained with hematoxylin and eosin (H&E) for histopathological evaluation by a board-certified pathologist specializing in Head and Neck Pathology. Histologic diagnosis, extent of tumor necrosis, mitotic activity, presence of keratinization and tumor grade (well, moderately and poorly differentiated) were analyzed.

### Short Tandem Repeat Testing

Short tandem repeat (STR) testing was performed to confirm the identity of PDXs. Briefly, DNA was prepared from fast frozen tissue or paraffin embedded tissue using Qiagen DNeasy Blood and Tissue kit (Catalog #69504). DNA was sent to Genetica Labcorp for STR testing and profiles compared between patient and PDX as previously described (6).

### RNA Sequencing

Total RNA was isolated and RNA sequencing libraries generated from flash frozen tissue as biologic triplicates by GeneWiz (South Plainfield, NJ). Final libraries were quantified on a high-sensitivity bioanalyzer chip and sequenced on the HiSeq (Illumina). Bulk RNA-Seq files are pre-processed for quality control using fastp with default parameters. STAR aligner in two-pass mode is used to align the reads to a combined murine and human reference genome: GRCh38 and GRCm38 (7).

### Statistical Analysis

To assess if there was a significant difference in fresh vs. cryopreserved implantation methods with respect to days-until-passage, a paired t-test was run at the patient level. To assess if there were differences in implantation methods with regards to keratin pearl, mitosis or necrosis, separate linear mixed models were fit in R (3.6.2), where patient was modeled as random effect. The resulting p-values were estimated using Satterthwaite’s approximation.

## Results

### Efficiency of PDX Tumor Establishment

Ten individual patient tissues were received from the Biobank. Tissues were split and implanted under two conditions: 1) fresh tissue procured and implanted within 48 hours of receipt; and, 2) frozen tissue, cryopreserved within 48 hours of receipt and implanted at a later date (at least 48 hours after cryopreservation). All PDX’s were generated from patients diagnosed with squamous cell carcinoma with the exception of UW-SCC-97 which was a metastatic melanoma as seen in Table 1.

**Table 1.** PDX establishment success by condition and patient characteristics.

| Patient | Implantation Conditions<br>(Growth Observed) |  | Patient Data |  |  |  |  |  |  |  |  |
| --- | --- | --- | --- | --- | --- | --- | --- | --- | --- | --- | --- |
|  | Fresh | Cryo-preserved | HPV Status | Primary tumor type | Origin of Tissue | Diagnosis | Sex | Age | Tobacco History | Metastatic | Original staging |
| UW-SCC-97 | Yes | Yes | - | Skin | Lymph node | Melanoma | F | 71 | No | Yes | NA |
| UW-SCC-100 | No | No | unknown | Tongue | Tongue | Squamous cell carcinoma | M | 50 | No | No | pT2N2b |
| UW-SCC-102 | No | No | + | Tongue | Tongue | Squamous cell carcinoma | M | 60 | Yes | No | pT2N1 |
| UW-SCC-103 | No | No | - | Floor of Mouth | Floor of Mouth | Squamous cell carcinoma | M | 57 | Yes | No | pT4aN0 |
| UW-SCC-130 | Yes | Yes | - | Skin | Parotid gland | Metastatic squamous cell carcinoma | M | 85 | No | Yes | TxN2bM0 |
| UW-SCC-133 | Yes | Yes | + | Base of Tongue | Tongue | Recurrent invasive squamous cell carcinoma | M | 69 | Yes | No | pT4NxMx |
| UW-SCC-136 | Yes | Yes | - | Hypopharynx | Hypopharynx | Squamous Cell Carcinoma | M | 59 | Yes | No | T2N1M0 |
| UW-SCC-137 | Yes | No* | - | Nasal Cavity | Ethmoid Sinus | Poorly Differentiated Carcinoma | M | 77 | No | No | pT4b |
| UW-SCC-148 | Yes | Yes | - | Unknown primary | Lymph Node | Poorly Differentiated Squamous Cell Carcinoma | M | 60 | Yes | Yes | TxN3M0 |
| UW-SCC-149 | Yes | Yes | - | Skin | Lymph node | Squamous cell carcinoma/spindle cell carcinoma | F | 71 | Yes | Yes | pT2NxMx |

\*No invasive SCC developed, cryopreserved tissue developed a lymphoproliferative mass

Overall, tumor growth was seen in 70% (95% CI: 35%-93%) of tissues. Three of the patient tissues, did not grow in either condition while the remaining 7 grew in at least one condition. One tumor (UW-SCC-137) grew a SCC from the fresh implant but a lymphoproliferative mass from the cryopreserved tissue. Another, UW-SCC-130, had two distinct masses develop from the cryopreserved specimen. One was a SCC with an appearance similar to the primary and to the fresh implantation specimen while the second demonstrated a lymphoproliferative process. The development of lymphoproliferative masses has been previously described (8,9).

### Characteristics of resulting PDXs

Tumors growing in mice were monitored at least weekly. Tumors were harvested when they reached a size of >500 mm^3^. We identified no difference between fresh injection and cryopreserved tumors in time from implantation to passage (Figure 1A, p=0.53).

**Figure 1.**
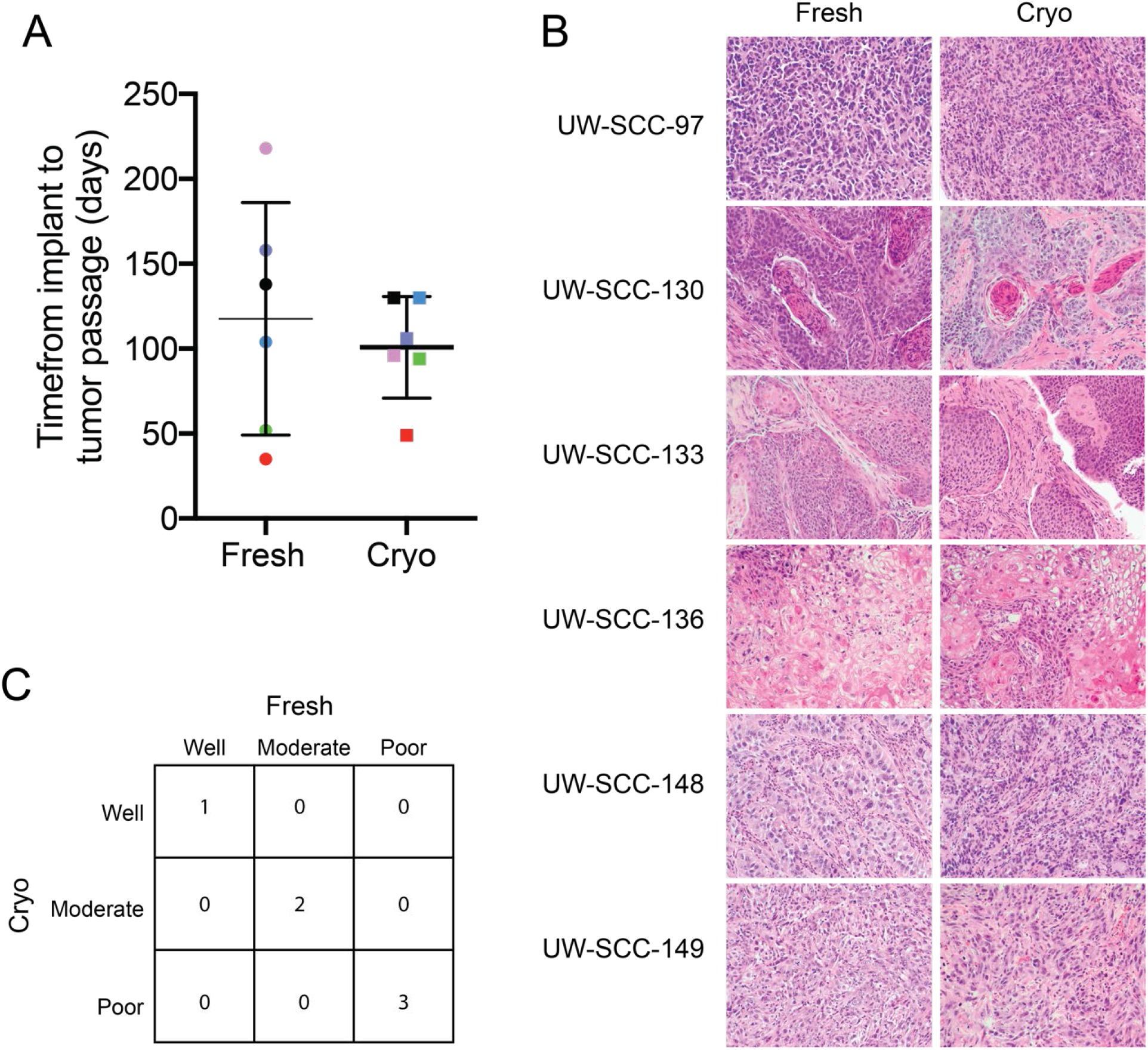
Comparison between fresh and cryopreserved implantation of tumor tissue. **A)** Time from tumor implantation to first passage (when tumor reached a size of 500mm^3^) was not different (p=0.53). Each color represents a different PDX. **B)** Hematoxylin and eosin stained paraffin sections (Magnification 10x). **C)** Matrix table comparing tumor grade of patient tissue between conditions, as scored by a pathologist.

H&E sections of the tumor (Figure 1B) were analyzed by a pathologist specializing in head and neck pathology who confirmed the histologic subtype. We assessed extent of tumor necrosis, presence of keratinization, and mitotic activity (Table 2). No significant difference in any of these parameters (p=0.12, 0.34 and 0.46, respectively) was identified between fresh and cryopreserved injection. Squamous cell carcinomas also had the tumor grade (well, moderately and poorly differentiated) analyzed (Figure 1C). No significant deviations were found in overall tissue morphology or tumor grade between tumors established by the two different methods.

**Table 2.**
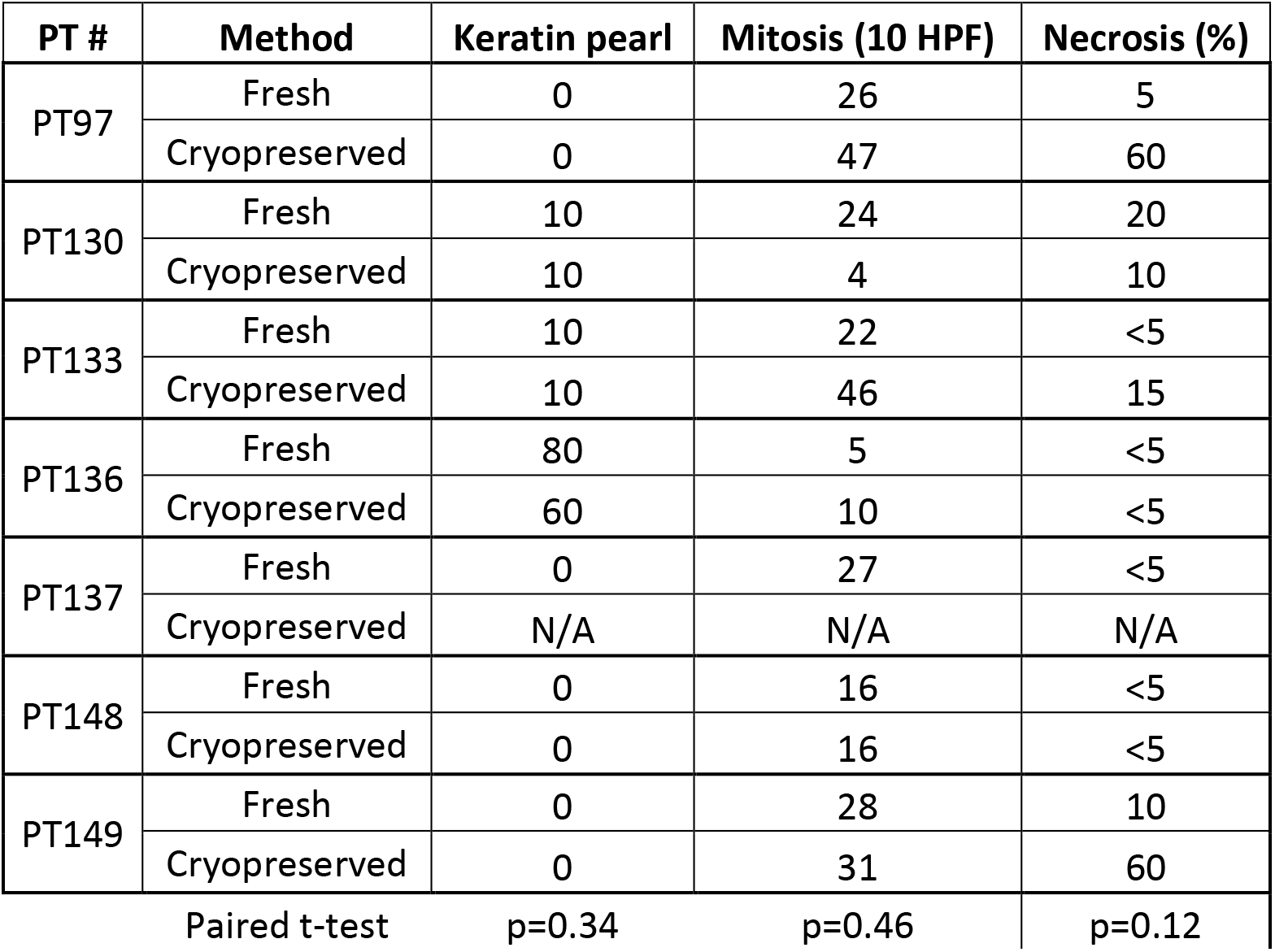
Comparison of histologic parameters between fresh and cryo-preserved PDXs.

PDXs for which original patient tissue and P1 tissue from both conditions could be procured were STR tested. A high degree of similarity was seen for all patient/PDX pairs (Table 3 and Supplemental Table 1).

**Table 3.** Stability of STR profile. Percent Match Algorithm showing percentage of allele match between original patient tissue and P1 of fresh and cryopreserved patient tissue. Samples with no growth excluded.

|  | Patient (Percent Match Algorithm %) |  |  |  |  |  |  |
| --- | --- | --- | --- | --- | --- | --- | --- |
|  | UW-SCC-97 | UW-SCC-130 | UW-SCC-133 | UW-SCC-136 | UW-SCC-137 | UW-SCC-148 | UW-SCC-149 |
| Fresh | 100 | 100 | 100 | 100** |  | 100 | 96 |
| Cryo-preserved | 100 | 100 | 100 | 100** | lymphoproliferative | *79 | 96 |

\* PT148 frozen tissue was from P2 rather than P1
\*\*No patient tissue was available for STR analysis so match shown is between Fresh and Cryopreserved samples.

RNA-Seq analysis was performed on Matched fresh and cryopreserved PDXs were analyzed by RNA-Seq analysis to explore differences in gene expression based on route of establishment. We aligned reads to a combined murine and human hybrid reference genome. Only a small proportion of reads mapped to the murine component and no difference was seen between fresh and cryopreserved samples. (Figure 2A). Multidimensional scaling (MDS) of matched fresh and cryopreserved PDXs revealed clustering of PDXs based on handling before implantation (Figure 2B). UW-SCC-130 showed good clustering within implantation (i.e. cryopreserved or fresh) and slight separation between approach. UW-SCC-136 also showed good clustering within implantation approach but demonstrated significant separation between components 1 and 2. To further investigate this difference, percentage of reads aligned to genes and exons was compared. While for most samples the majority of reads showed good coverage of genes and exons, UW-SCC-136 was a significant outlier within the fresh tissue samples (Supplemental Figure 1).

**Figure 2.**
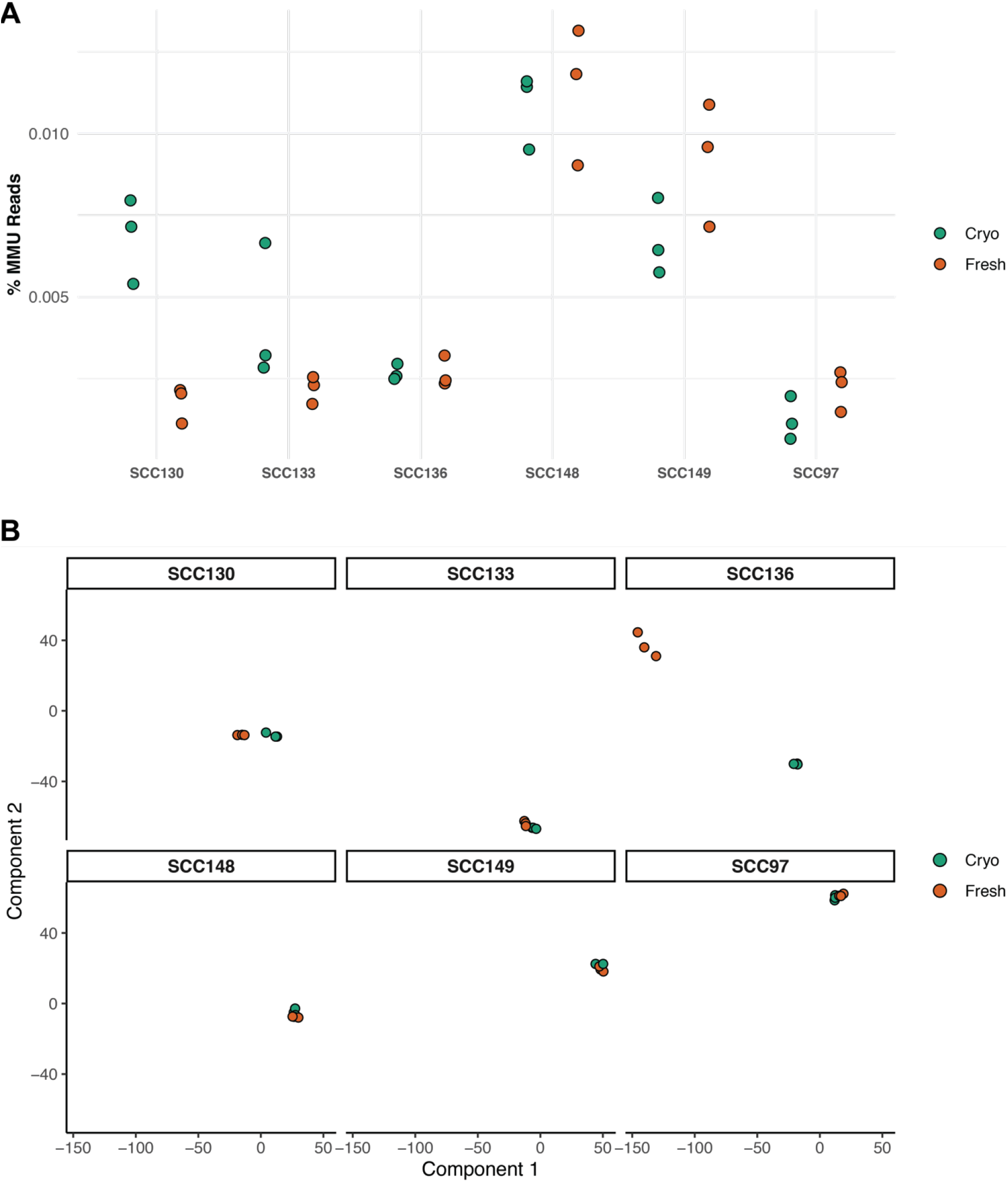
RNA-Seq analysis of fresh and cryopreserved specimens. **A)** No difference in the proportion of reads mapping to the murine genome was seen based on method of implantation. Three separate tumor specimens from each condition underwent RNA-Seq analysis. **B)** Multidimensional scaling (MDS) clustering of fresh and cryopreserved PDX tissue demonstrates similarity for most PDXs.

## Discussion

PDX’s have become an integral part of oncology research and are currently used across the spectrum of cancer research ranging from drug development to biomarker analysis (3–5, 10–16). While the use of PDX’s is expected to continue and expand as PDX lines become further annotated and characterized, the generation, characterization and maintenance of PDX’s requires a significant investment of time and resources. Herein, we explored the feasibility of cryopreserving patient tissue prior to implantation. The ultimate goal is to conserve precious resources by allowing tumors to be characterized prior to generating the PDX.

The acquisition of patient tissue within an academic institution involves a highly coordinated effort that begins from the time a patient is diagnosed and extends until after a PDX is established from remnant tissue. At our institution, the workflow begins with the physician or staff identifying a case suitable for donation of remnant cancer tissue. The patient is consented to an IRB approved protocol by a research team member. The surgical team is notified, patient tissue is collected in saline (not formalin) and given to surgical pathology. Pathology provides confirmation of diagnosis and staging while also allocating tissue to the biobank and research teams. The tissue is retrieved from the biobank and processed for implantation and/or cryopreservation. We have previously shown that tumor should be processed within 48 hours of harvest to maximize the efficiency of establishing xenografts (5). Mice are regularly observed for tumor growth and animal health and when a tumor develops, it must be passaged and/or cryopreserved. Simultaneously, various analyses can be performed on remnant tissue to help the investigator identify the optimal PDX for a specific question. These studies can include simple histologic analysis, immunohistochemistry, in situ hybridization, targeted or untargeted gene sequencing, among other tests. By immediately cryopreserving samples these analyses can be performed prior to implantation of the tumor thus saving the resources that would otherwise be used to establish PDXs which may never be used experimentally.

We saw no negative impact of an immediate cryopreservation approach. For most PDXs, tumors developed from both fresh and cryopreserved tissues. Lymphoproliferative masses arose infrequently, but from both fresh implantation and cryopreserved tumors. The overall histology did not otherwise differ between approaches and the time from implantation to passage demonstrated greater patient-to-patient heterogeneity than approach-based differences.

We acknowledge clear limitations to this work. Most notably, due to the interests of our lab, we utilized only samples from patients with malignancies of the head and neck. Whether our results translate to other malignancies requires further study. Hernandez and colleagues assessed a similar approach in hepatobiliary and pancreatic cancers and demonstrated similar findings (17) suggesting that this approach is likely to work in a wide variety of cancer types. In addition, while we have focused on the potential benefits of cryopreserving samples prior to implantation, we acknowledge that this requires us to perform our characterization on a cohort of patients whose tumors may never establish PDXs. If characterization involves clinically appropriate testing, costs for this might be defrayed. However, for those cases requiring non-clinical analysis, some percentage of samples may be analyzed and ultimately be unable to be reanimated into PDXs.

In conclusion, we have demonstrated that it is feasible to cryopreserve fresh patient tissue to be used at a later timepoint to generate PDXs. This provides an option by which investigators can establish large libraries of potential PDXs which can be animated at the time they are needed rather than only when they are obtained. This may allow investigators to more quickly establish diverse and comprehensive libraries, encourage resource sharing, and accelerate discovery.

## Declarations

Ethics approval and consent to participate – the generation of patient derived xenografts from deidentified tissues has been deemed to be not-human subjects research and exempt from IRB review. Animal work has been approved by the University of Wisconsin Institutional Animal Care and Use Committee under an approved animal protocol.

### Consent for publication

All authors have approved the final article and consent to publication.

### Availability of data and materials

The datasets used and/or analyzed during the current study are available from the corresponding author on reasonable request.

### Authors’ contributions

LA, AD, RH, CL, AM analyzed and interpreted data. RH performed histologic analysis. CL performed statistical analyses. DW, JGS, and RJK provided concept, oversaw data analysis, and writing of the manuscript. All authors have approved the final article.

### Acknowledgements

none

---

**Supplemental Figure 1.**
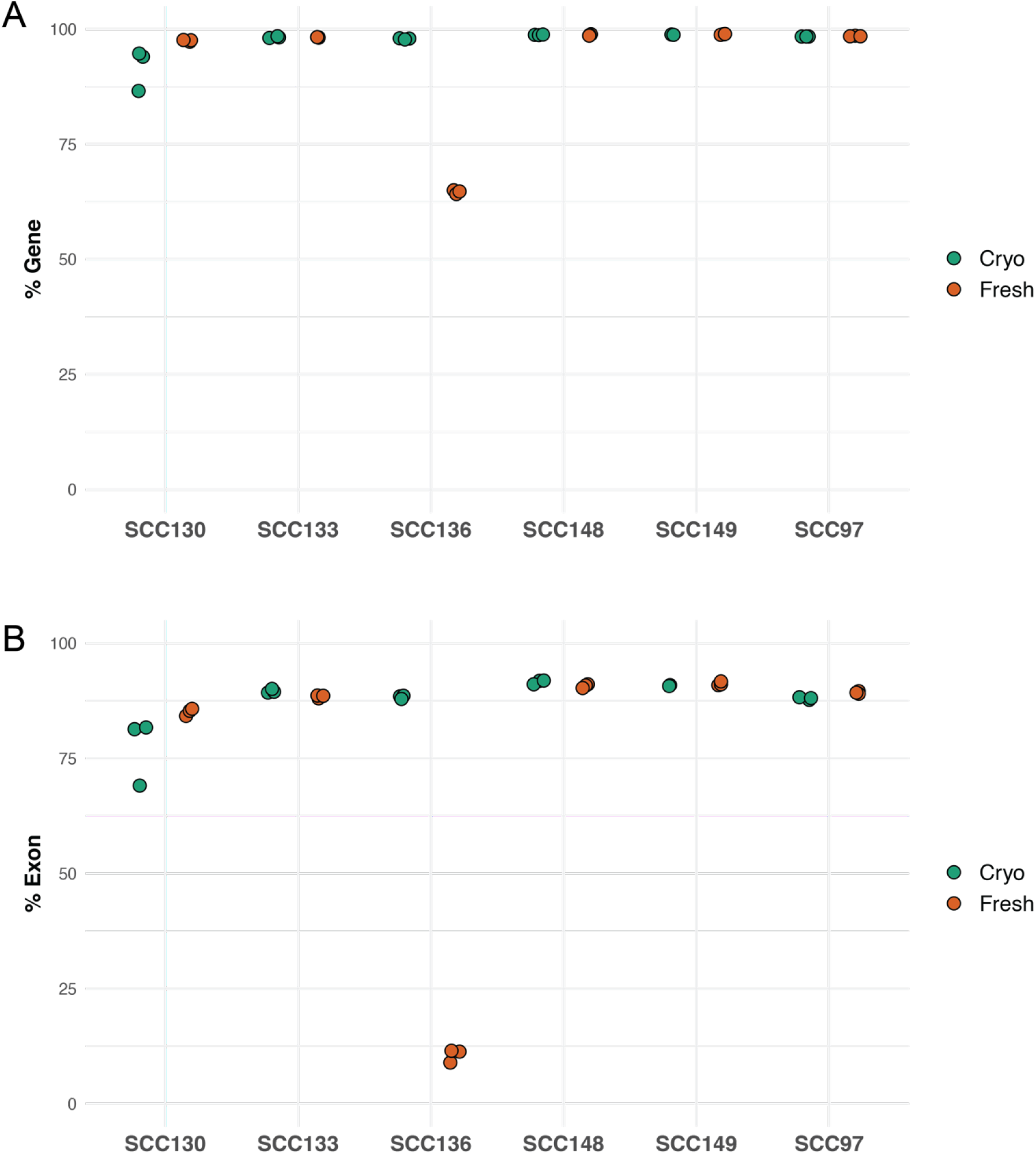
Alignment of samples to genes (A) and exons (B) demonstrates poor alignment of the fresh samples from UW-SCC-136 suggestive of possible DNA contamination.

**Supplementary Table 1.**
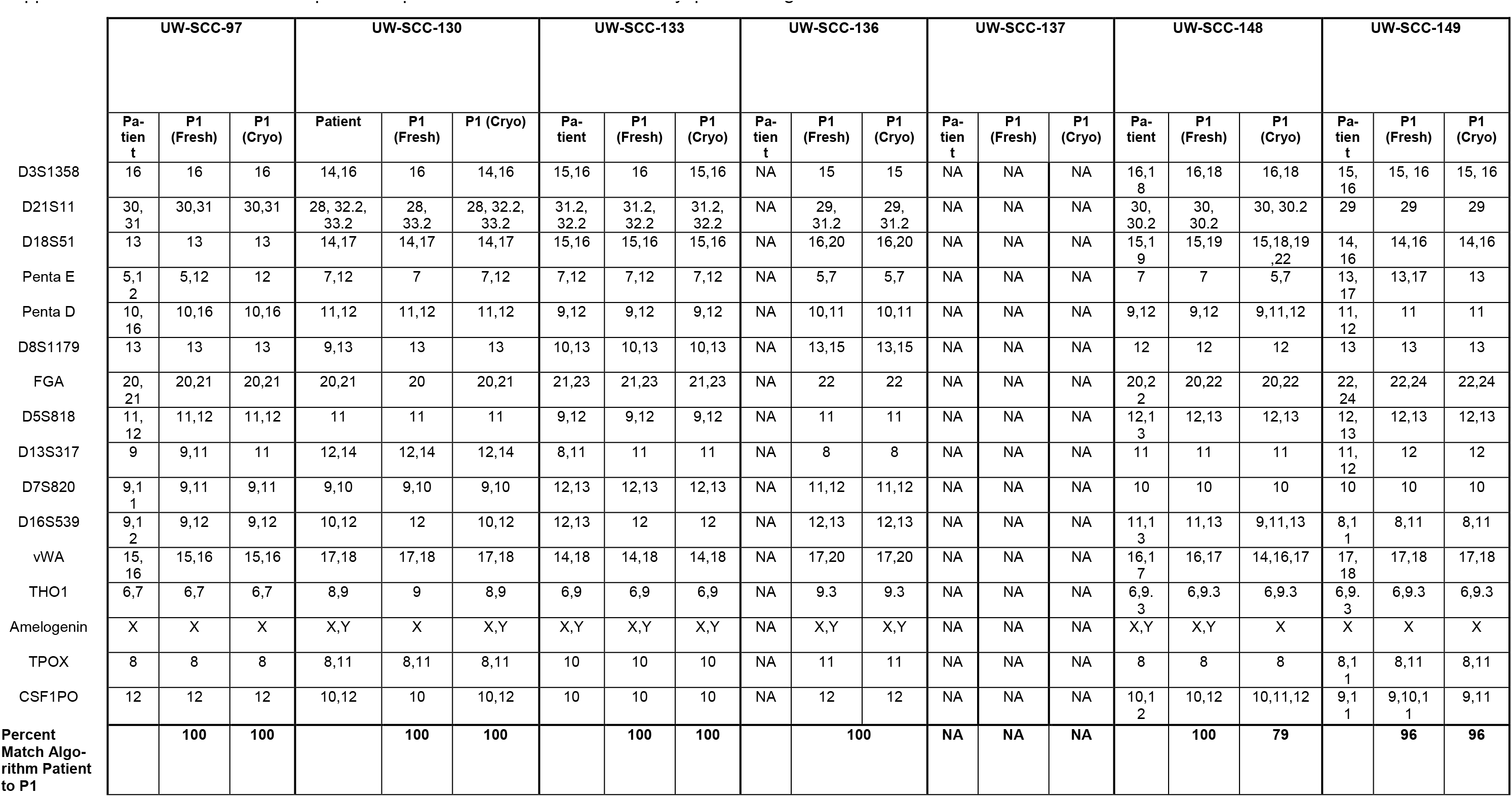
Detailed STR profiles of patient and PDX from fresh or cryopreserved growth.

